# Glycation proteomics identifies creatine kinase as a mediator of the effects of methylglyoxal on mitochondrial permeability transition

**DOI:** 10.64898/2026.10.01.756032

**Authors:** Rahiim F Lake, Sergiy M. Nadtochiy, Paul S. Brookes

## Abstract

Opening of the mitochondrial permeability transition (PT) pore is an important step in necrotic cell death in various pathologic conditions such as ischemia-reperfusion injury. Methylglyoxal (MGO) is a byproduct of glycolysis that covalently modifies proteins (i.e., glycation) and is widely regarded as toxic (glycative stress), contributing to the pathology of diabetes. Paradoxically, MGO has been shown to inhibit PT pore opening, but the mechanism of this effect is poorly understood. Herein we show that glycation is elevated in various cardioprotective scenarios. Furthermore, building on a serendipitous finding of the ability of phosphate to block MGO’s effect on the PT pore, we used proteomics to map potential sites mediating this effect. From this analysis, mitochondrial creatine kinase (CK) emerged as a phosphate-sensitive MGO target with links to the PT pore. Although MGO inhibited CK activity, this did not appear to be important for its effect on the PT pore. Rather, MGO modified sites on CK that are important for its membrane binding, which likely mediates the combined effects of MGO and phosphate on the PT pore. Furthermore, a consensus sequence for MGO modifications in mitochondria is reported.

**Graphical Abstract:** 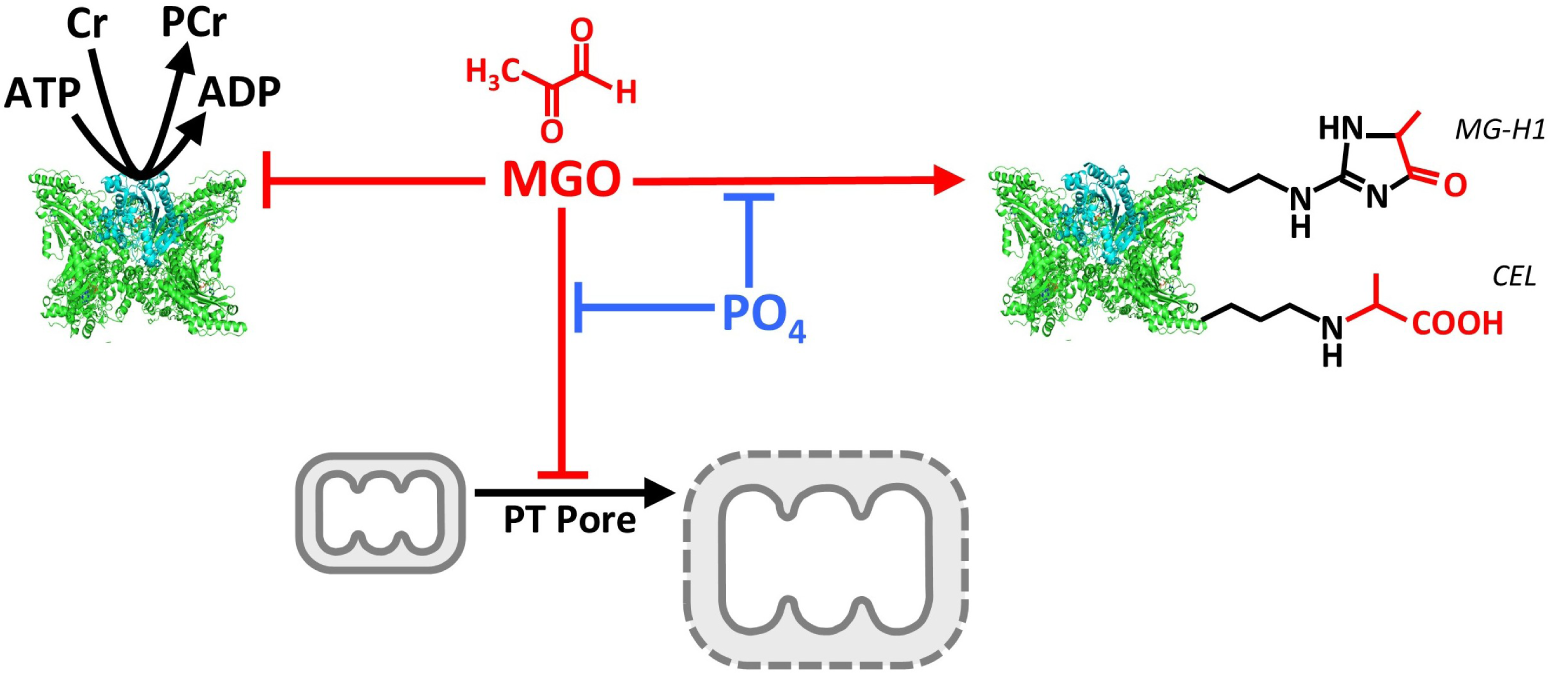

## Introduction

Blockage of coronary arteries starves cardiomyocytes of oxygen and substrates (aka ischemia), driving an increased reliance on glycolysis to make ATP, resulting in cytosolic acidosis. Na^+^/H^+^ exchange activity subsequently drives a cellular Na^+^ overload, followed by a Na^+^/Ca^2+^ exchange (NCX) driven Ca^2+^ overload ^[1]^. This Ca^2+^ enters mitochondria, and upon tissue reperfusion, mitochondrial Ca^2+^ overload is accompanied by a burst of reactive oxygen species (ROS) generation, activating the mitochondrial permeability transition (PT) pore, leading to cell death ^[2]^.

In diabetes, excessive glucose can post-translationally modify hemoglobin, generating the diabetic biomarker HbA1c in a process known as glycative stress. Furthermore, the glycolytic intermediate dihydroxyacetone phosphate (DHAP) can spontaneously degrade into the reactive glycating agent methylglyoxal (MGO) ^[3,4]^. MGO is generally regarded as toxic, and is scavenged by glutathione and the glyoxalase enzymes (Glo1 and Glo2) which detoxify MGO into d-lactate ^[3–5]^. Unchecked, MGO’s reaction with nucleophilic amino acid residues such as lysine and arginine drives the production of advanced glycation-end products (AGEs) including *N*-ε-carboxyethyllysine (CEL) and methylglyoxal-derived arginine hydroimidazoles (MG-H 1/2/3), as well as protein crosslinks ^[3,4,6]^. Although glycolytic MGO is made in the cytosol, it can permeate cell membranes ^[7]^. In this framework, glycolysis in ischemia-reperfusion (IR) injury should be viewed as an initiator of pathology.

The focus of the glycative stress field on diabetes and cardiovascular pathology has led to the potentially beneficial signaling functions of MGO being largely overlooked. Although the vascular pathologic effects of MGO may increase the risk or incidence of cardiac syndromes such as heart-attack, many studies show an association between accelerated glycolysis or increasing glycative stress, and a reduction in damage from heart attack. A key example of this is the finding that type-1 diabetic hearts are protected against acute IR injury ^[8–10]^. In addition, human epidemiology studies find no association between MGO levels and cardiovascular disease (CVD) in pre-diabetic patients ^[11]^. Human studies also find an association between high dietary MGO intake and lower inflammation ^[12]^, greater insulin sensitivity, and a lower prevalence of type-2 diabetes ^[13]^. Likewise in mice high circulating MGO levels have no effect on the cerebral vasculature ^[14]^. In humans, MGO scavengers can lower glycation markers (AGEs), but this does not impact vascular function ^[15]^. Furthermore, MGO is known to activate the protective Nrf-2 antioxidant signaling pathway ^[16,17]^.

Regarding cardiac IR injury, an association between elevated glycolysis and cardioprotection is seen in ischemic preconditioning (IPC) ^[18,19]^, the Glyco-Hi mouse ^[20]^, glucose insulin potassium (GIK) treatment ^[21–24]^, the *Alkbh7*^-/-^ mouse ^[25]^, SIRT1 activation ^[18,19]^, NAD^+^ supplementation ^[26]^, and insulin-Akt signaling ^[27,28]^. Notably, in-vitro studies have also shown a protective effect of the related molecule glyoxal in human trabecular muscles exposed to IR injury ^[29]^. Elevated DHAP (the precursor for MGO) is found in both cardiac and neuronal IPC ^[30,31]^, and elevated glyoxal/dicarbonyl metabolism is seen in serum of hypoxically preconditioned athletes ^[32]^ and mice ^[33]^.

Given the potential link between glycolysis, MGO and <u>protection</u> from IR injury, it is not surprising that MGO and phenylglyoxal (PGO) have previously been shown to inhibit the mitochondrial PT pore ^[34–37]^. However, the molecular mechanism underlying this effect is unclear. In yeast mitochondria, a role for R_107_ on ATP-synthase subunit g in PGO-sensitivity was proposed ^[35]^, but it remains unclear if this applies to MGO or to mammalian mitochondria, and whether other proteins in the mitochondrion are targets for MGO.

The composition of the mitochondrial PT pore has been the subject of considerable debate ^[38]^. Classical studies proposed a multi-component structure spanning the mitochondrial inner and outer membranes and including the adenine nucleotide translocase (ANT), the voltage dependent anion channel (VDAC), the prolyl-isomerase cyclophilin D (CypD), the mitochondrial peripheral benzodiazepine receptor (PBR), and creatine kinase (CK). Initial knockout mouse studies suggested no role for ANT ^[39]^, and subsequent discoveries led to the proposal that the core component of the PT pore is ATP synthase (complex V) ^[40]^. A more recent consensus is that both ATP synthase and ANT plus the mitochondrial phosphate carrier (PiC) may all play a role in PT pore formation ^[41]^. There is also broad agreement on the role of CypD as a pore regulator, via binding to the OSCP subunit of ATP synthase ^[42]^. CypD is the target of the PT pore inhibitor cyclosporin A (CsA), which is cardioprotective in animal models of IR injury ^[43]^, although this has not translated into successful cardioprotection in human clinical trials ^[44]^. Despite the growing consensus on ATP synthase and CypD as PT pore components ^[38]^, the role of other peripheral proteins such as CK has received less attention.

Herein, we performed a proteomics analysis to identify MGO targets in mitochondria, which identified several proteins implicated in the PT pore. Serendipitously during these studies we discovered that the effect of MGO on the PT pore was lost in the presence of inorganic phosphate. Subsequent studies revealed differential MGO modification of several proteins in the presence or absence of phosphate, including S-type (mitochondrial) creatine kinase. Herein, the interactions between MGO, CK and the PT pore are explored in more detail.

## Materials & Methods

### MGO, Quantitation & Phosphate Reactivity

MGO was distilled from commercial stock solutions (∼40% by volume) ^[7]^. The concentration of distilled MGO was determined by reaction with O-phenylenediamine (OPD), monitoring absorbance of the 2-methylquinoxaline product by spectrophotometry at 320 nm ^[45]^. To determine if phosphate was reactive toward MGO, the OPD assay was repeated with pre-incubation of 1 mM MGO and 3 mM phosphate. This resulted in only a ∼1.2% difference in the yield of 2-methylquinoxaline, indicating no reactivity between phosphate and MGO.

### Animals

Mice were on the C57BL/6J background, maintained on a 12h light/dark cycle with food and water available *ad libitum*, in a pathogen-free and AAALAC accredited vivarium (animal protocol number: UCAR-2007-087). Both sexes were used. Mice were anesthetized with tribromoethanol (Avertin, 200 mg/kg IP) prior to experiments. Samples of heart tissue from wild-type and Glyco-Hi mice ^[20]^ were generously provided by Dr. Bradford Hill (University of Louisville).

### Perfused Hearts

Mouse hearts were retrograde/Langendorff perfused as previously described ^[46]^, in Krebs-Henseleit buffer (KHB) comprising 118 mM NaCl, 4.7 mM KCl, 1.2 mM MgSO_4_, 25 mM NaHCO_3_, 1.2 mM KH_2_PO_4_, 2.5 mM CaCl_2_, 5 mM glucose, 0.2 mM pyruvate, 1.2 mM lactate, and 100 μM palmitate conjugated 6:1 with fat-free BSA. KHB was gassed with 95% O_2_ and 5% CO_2_ at 37°C to maintain pH 7.4. After stabilization, 1 mM MGO was introduced to perfusion media via a port immediately above the perfusion cannula, for a period of 5 min. Hearts were then snap-frozen and pulverized in liquid N_2_, then homogenized in 1 ml of 1 mM aminoguanidine. 100 μl aliquots were stored at −80°C for proteomics analysis. Ischemic preconditioning was performed as previously ^[19]^, with 3 cycles of 5 min. ischemia plus 5 min. reperfusion. For modeling of GIK ^[21–24]^, hearts were perfused for 20 min. with regular KHB (5mM glucose, 100 μM palmitate-BSA) or supplemented with glucose (12 mM total) plus insulin (1 mU/ml).

### Isolated Mitochondria

Mouse liver tissue was diced with scissors in ice-cold liver mitochondrial isolation medium (LMIM, 250 mM sucrose, 10 mM Tris, 1 mM EGTA, pH 7.3 @ 4°C), rinsed to remove blood, then homogenized with 10 strokes in a glass Dounce homogenizer. Homogenate was centrifuged at 800 x *g* for 3 minutes, then the supernatant centrifuged at 10,000 x *g* for 10 minutes. The pellet was subjected to 2 further centrifugation wash steps and finally suspended in 0.5 ml of LMIM. Mouse heart tissue was chopped in heart mitochondrial isolation media (HMIM, 300 mM sucrose, 10 mM Tris, 1 mM EGTA, pH 7.3 @ 4°C) and homogenized with 25 x 1 s. passes in a Tissumizer (IKA Inc.) at 6000 rpm. Homogenate was centrifuged at 800 x *g* for 5 minutes, and supernatant was passed through a 300 μm mesh then centrifuged at 7000 x *g* for 10 minutes. The pellet was subjected to 2 further centrifugation wash steps and resuspended in 0.5 ml of HMIM. Mitochondrial protein content was assayed by the Folin-Ciocolteau reagent ^[47]^.

### Mitochondrial Permeability Transition Pore Opening

Mitochondrial permeability transition (PT) pore opening was monitored spectrophotometrically at 540 nm, as Ca^2+^ induced swelling/light-scatter. Mitochondria were incubated at 0.5 mg protein/ml., with succinate as respiratory substrate. The known pore inhibitor cyclosporin A (CsA, 0.2 μM) was used as a control. Where present, MGO was added to the mitochondrial suspension and incubated for 5 min. prior to addition of Ca^2+^. Based on our previous studies, initially liver mitochondria were studied in a swelling buffer comprising 25 mM sucrose, 195 mM mannitol, 40 mM HEPES, pH 7.3 at 37°C ^[48]^, while heart mitochondria were studied in a swelling buffer comprising 120 mM KCl, 50 mM Tris, 3mM KH_2_PO_4_, pH 7.3 at 37°C ^[49,50]^. This difference in buffer composition led to the discovery that phosphate inhibits the effect of MGO on the PT pore. As such, subsequent studies on heart mitochondria were performed in the liver-specific buffer, initially without phosphate. Four conditions were tested: (i) 100 μM CaCl_2_. (ii) 100 μM CaCl_2_ + 1 mM MGO. (iii) 100 μM CaCl_2_ + 3 mM phosphate. (iv) 100 μM CaCl_2_ + 3 mM phosphate + 1 mM MGO.

### Proteomics

Mitochondria were suspended in PT pore buffer (see above) at 0.5 mg protein/ml., with succinate as respiratory substrate. MGO and/or phosphate were added as above, without Ca^2+^ to initiate swelling. After 5 min., 1 mM aminoguanidine was added to quench excess unreacted MGO, and mitochondria were pelleted by centrifugation then snap frozen. Heart or mitochondrial samples were processed for proteomics (Trypsin digest, desalting etc.) in the URMC proteomics core, and analyzed for posttranslational modifications. Specific modifications searched were CEL (+72.021129 Da) and MG-H1 (+54.010565 Da). Flanking amino acid sequences around glycation sites were obtained using a custom Python script and Uniprot accession numbers.

### Creatine Kinase Assay

CK enzyme activity was measured spectrophotometrically in a system with accessory enzymes pyruvate kinase and lactate dehydrogenase (PK/LDH mix, Sigma #P0294), in a buffer comprising 150 mM KCl, 100 mM triethanolamine, 10 mM EDTA, 16 mM MgSO_4,_ pH 7.6 at 37°C. Heart mitochondria (25 μg) were incubated for 30 minutes in 700 μl of assay buffer containing varying concentrations of MGO (0 – 750 μM) prior to being assayed for CK activity. Where applicable, 3 mM KH_2_PO_4_ was present in buffers prior to MGO addition. To each cuvet was then added 2 μl of PK/LDH mix, 0.9 mM phosphoenolpyruvate, 4 mM ATP, 1 mM NADH, 10 μM rotenone, and 10 μg/ml oligomycin, all in 100 μl of a working stock solution. The assay was started by adding 200 μl of 120 mM creatine monohydrate, bringing the final volume to 1 ml. Disappearance of NADH was monitored at 340 nm.

## Results

The complete original data set for this paper is stored on the data sharing site FigShare, at DOI: 10.6084/m9.figshare.34047288

### Cardioprotection Associated with Elevated Glycation

Swe previously showed that hearts from *Alkbh7*^-/-^ mice, which are protected against IR injury, exhibit elevated levels of MGO protein adducts, measured by western blot ^[25]^. In a similar manner, **Suppl. Fig. 1.** shows that hearts subjected to IPC also exhibit more glycation, as do those subjected to GIK therapy. In addition, hearts from mice with a deletion of PFK-2, which results in faster glycolysis (aka Glyco-Hi mice ^[20]^) also showed more MGO protein adducts. As such, four separate models of cardioprotection all exhibit elevated protein glycation, questioning the narrative that MGO is purely pathologic.

### Glycation Proteomics of the Intact Mouse Heart

Mouse hearts were treated with 1 mM MGO for 5 min, homogenized in the presence of aminoguanidine to quench unreacted MGO, then subjected to proteomics analysis to detect peptides containing MGO-modified lysine (CEL) or arginine (MG-H1) (**Fig. 1A**). Pathway analysis (ShinyGO ^[51]^) revealed enrichment in mitochondrial and metabolic pathways, and cardiac contractile proteins (**Fig. 1B**), the latter being consistent with previous findings of extensive glycation in the cardiac contractile machinery ^[52,53]^.

**Figure 1.**
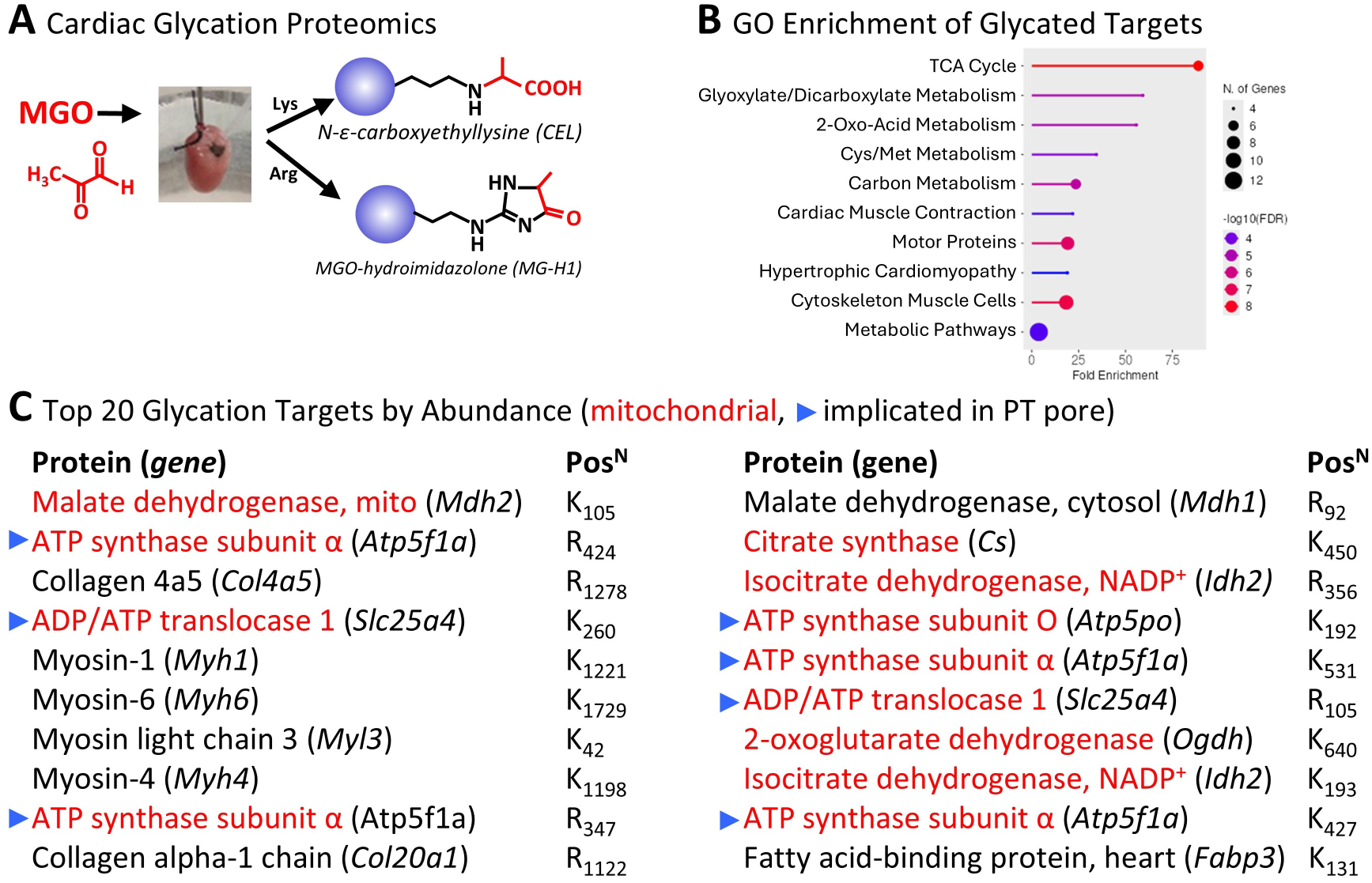
Cardiac Glycation Proteomics. **(A)**: Mouse hearts were Langendorff perfused and treated for 5 min. with 1 mM MGO, then snap frozen and processed for proteomics analysis to identify peptides modified with N-ε-carboxyethyllysine (CEL) or arginine MGO-hydroimidazolone (MG-H1). All data are from N=4 animals per group, and the complete dataset is available on FigShare. **(B):** Pathway Analysis (ShinyGO ^[51]^) showing KEGG enriched pathways for MGO targets. **(C):** Top 20 glycation targets by abundance. The position of each modification within the protein sequence is listed. Hits in red are mitochondrial proteins, and the blue arrow denotes proteins implicated in mitochondrial PT pore opening.

A total of 66 target sites were identified across 52 proteins, with a roughly equal distribution between arginine and lysine residues. Notably, of the top 20 most abundant modified peptides, 12 were mitochondrial, and 7 of these have previously been implicated in the mitochondrial PT pore, including several subunits of ATP synthase, as well as adenine nucleotide translocase (ANT) (**Fig. 1C**). Although glycolytic MGO is synthesized in the cytosol, these data clearly indicate that MGO can cross the mitochondrial membrane(s) to modify proteins within the organelle.

### Glycation Proteomics of Cardiac Mitochondria

To investigate the process of mitochondrial protein glycation more thoroughly, isolated heart mitochondria were incubated with MGO, followed by proteomic analysis to detect peptides containing MGO-modified lysine or arginine (**Fig. 2A**). A total of 176 modifications were identified on 55 individual proteins, including many of the same PT pore components identified at the whole heart level (**Fig. 2B**). Notably, unlike whole heart tissue, mitochondrial MGO modified proteins were skewed toward arginine (113) vs. lysine (63). This prompted us to analyze the sequences surrounding the modified residues, to determine if a consensus sequence for MGO modification could be elucidated.

**Figure 2.**
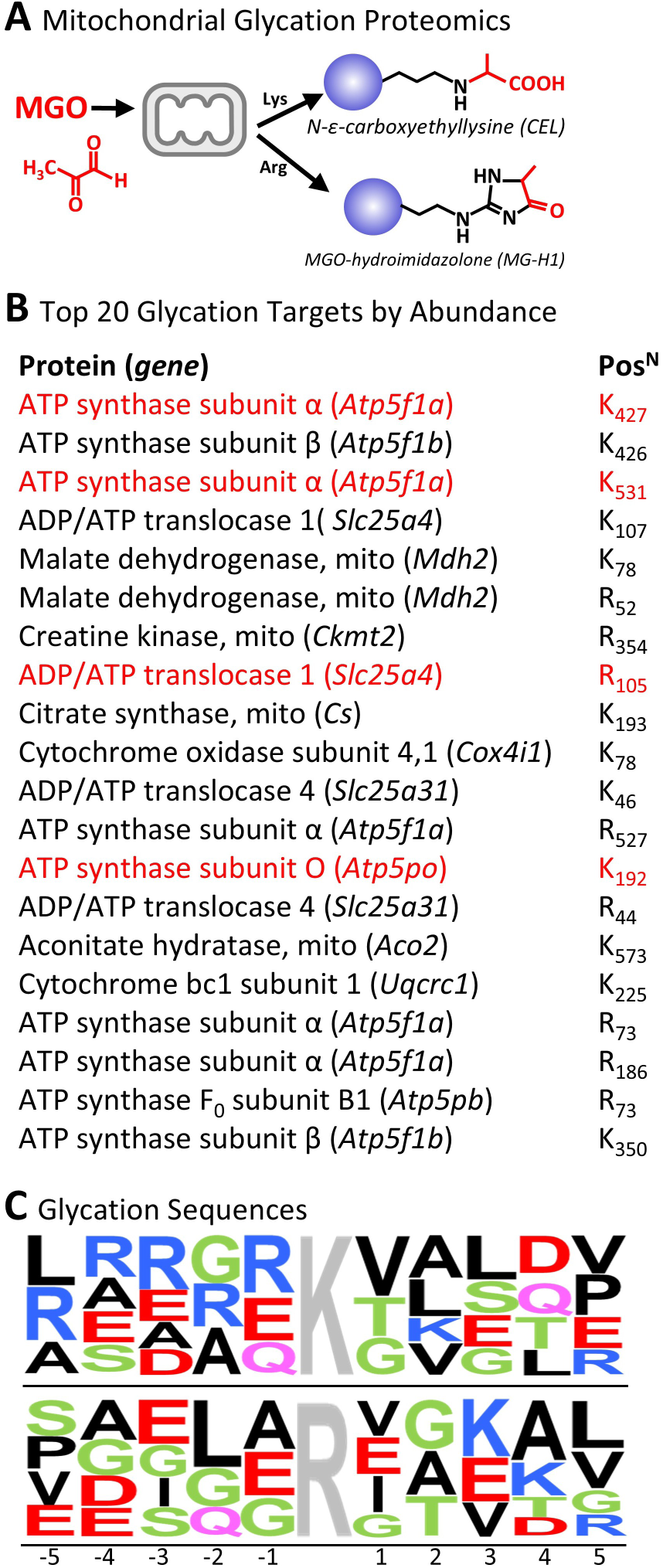
Cardiac Mitochondrial Glycation Proteomics. **(A)**: Mouse heart mitochondria were treated for 5 min. with 1 mM MGO, then snap frozen and processed for proteomics analysis to identify peptides modified with N-ε-carboxyethyllysine (CEL) or arginine MGO-hydroimidazolone (MG-H1). All data are from N=5 independent mitochondrial preparations per group, and the complete dataset is available on FigShare. **(B):** Top 20 glycation targets by abundance. The position of each modification within the protein sequence is listed. Hits in red are the same as identified in the whole heart glycation proteome in Fig. 1. **(C):** Weblogo diagrams showing the relative abundance of flanking amino acids in glycated peptides. For visual clarity, only residues that attained >8% abundance in a given position are shown.

As shown in the WebLogo diagrams (**Fig. 2C**), this analysis revealed that for modified lysine, the presence of an upstream R or E and downstream small aliphatic residues (VAL) was predictive for glycation. Meanwhile for modified arginine, a prevalence of glycine residues was seen upstream, with similar small aliphatic residues downstream. Notably, methionine was rare (<3% abundance) and cysteine was completely absent near glycation sites, consistent with thiol residues as alternate electrophilic targets for MGO reactivity (i.e., these residues were not seen near modified K or R because they would have been modified instead of K or R). A previous analysis of actin and myosin glycation ^[53]^ revealed a slight favoring of leucine or glutamate surrounding the modified K or R, but this may be influenced by the focus on myofilament proteins with limited sequence variability. It is also possible the environment inside the mitochondrion (e.g. alkaline pH) may influence the sequence specificity of glycation reactions relative to that in the cytosol.

### MGO Inhibition of the PT Pore, Impact of Phosphate

We and others previously showed that MGO can inhibit Ca^2+^-induced opening of the mitochondrial PT pore ^[34–37]^. During a series of experiments examining differences between heart and liver mitochondria, we serendipitously discovered that the presence of phosphate in the buffer was able to prevent the ability of MGO to block PT pore opening. As shown in **Fig. 3A**, the presence of phosphate alone at physiologically relevant levels (3 mM) had a minimal effect on Ca^2+^ induced swelling, but it completely blocked the inhibitory effects of MGO on swelling.

**Figure 3.**
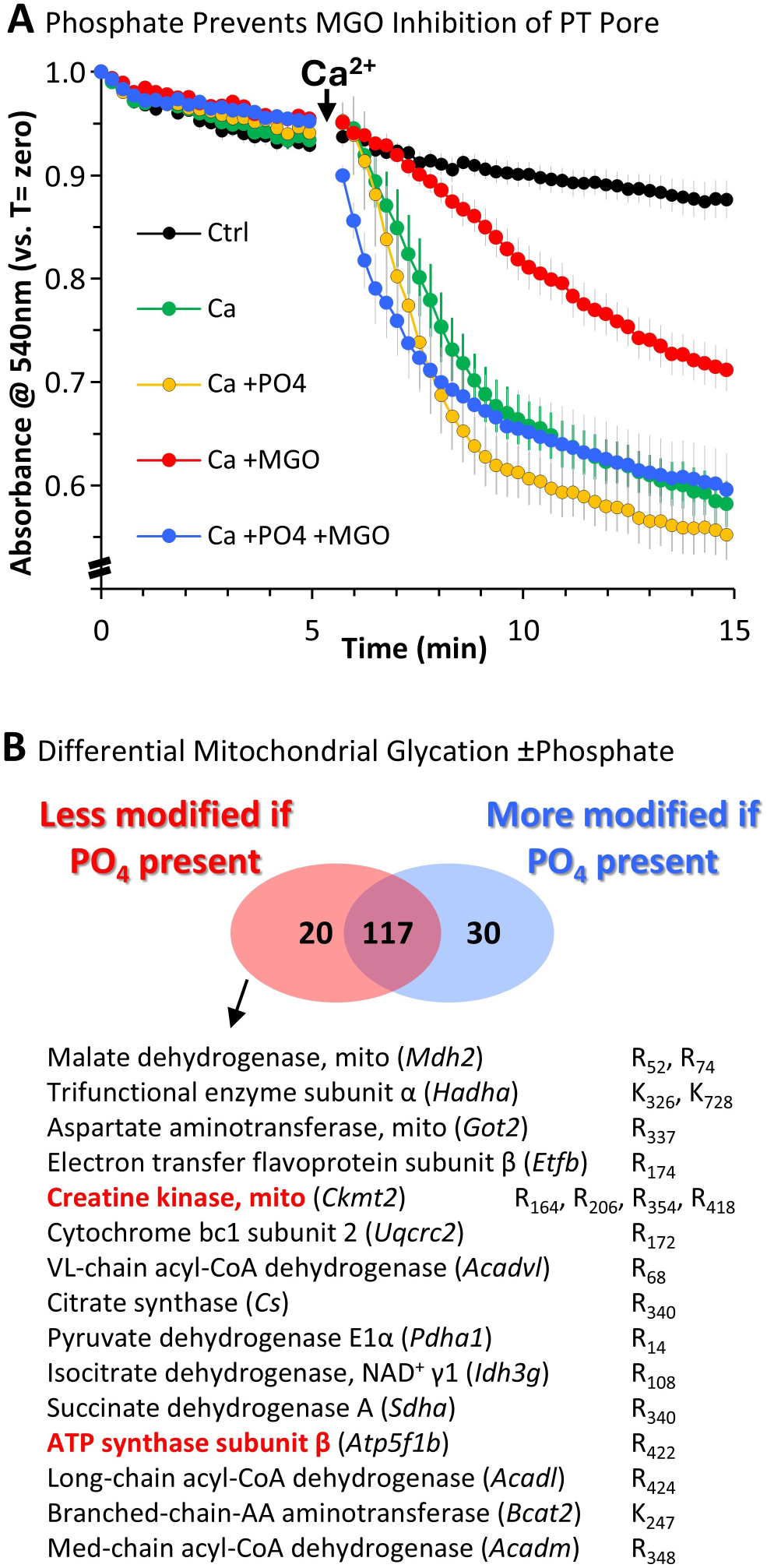
Phosphate and MGO Inhibition of the PT Pore. **(A)**: Mitochondrial PT pore opening was measured as spectrophotometric absorbance at 540 nm, corresponding to swelling-induced light scatter. 100 μM CaCl_2_ was added as shown by the arrow, and swelling was sensitive to the PT pore inhibitor CsA (not shown, but present in complete data set on Figshare). Where indicated, mitochondria were incubated with MGO (1 mM) and/or phosphate (3 mM). Data are means ± SEM, N=7-10 independent mitochondrial preparations. **(B):** Differential glycation proteomics of mitochondria treated with MGO ± phosphate as in panel A. Table below shows 20 modified peptides that decreased in abundance by >1.5-fold when phosphate was present. The corresponding 30 proteins that became more glycated when phosphate was present are shown in Suppl. Fig. 1. Proteomic data are from N=2 independent mitochondrial preparations.

### Impact of Phosphate in the Glycation Proteome of Heart Mitochondria

These PT pore observations prompted us to conduct a proteomic analysis to determine if any of the previously identified MGO target proteins in mitochondria were differentially modified in the presence or absence of phosphate. Similar to the experiments in **Fig. 2**, a set of 167 MGO modifications were found in 46 separate proteins. The majority of these (117) occurred regardless the presence of phosphate. However, as shown in **Fig. 3B**, 20 proteins became less glycated when phosphate was present, while 30 proteins became more glycated (**Suppl. Fig. 2**). Given the ability of phosphate to block PT pore inhibition by MGO, we focused on those proteins that were less glycated with phosphate present. Among these targets were several proteins implicated in the PT pore, including 4 sites on mitochondrial creatine kinase (*Ckmt2*).

### MGO Inhibition of Creatine Kinase, the Impact of Phosphate

**Fig. 4A** shows the crystal structure of human mitochondrial CKMT2 (P17540, pdb-4z9m). No crystal structure is available for mouse CKMT2 (Q6P8J7), but the human/mouse isoforms exhibit 96% sequence homology (**Suppl. Fig. 3**), so we examined the AlphaFold predicted structure instead. As **Fig. 4A** shows, two of the phosphate-sensitive MGO modifications (R_164_ and R_354_) were close to the ATP binding site and appear to coordinate with ATP. Consistent with this, MGO inhibited the enzymatic activity of purified CK (**Fig. 4B**) and this inhibition was also seen in intact mitochondria (**Fig. 4C**). Notably however, the presence of phosphate did not appear to impact the inhibition of CK by MGO (**Fig. 4D**). Although there was a small inhibitory effect of phosphate alone on CK activity (consistent with previous observations ^[54]^), the % inhibition by MGO was the same regardless of phosphate presence.

**Figure 4.**
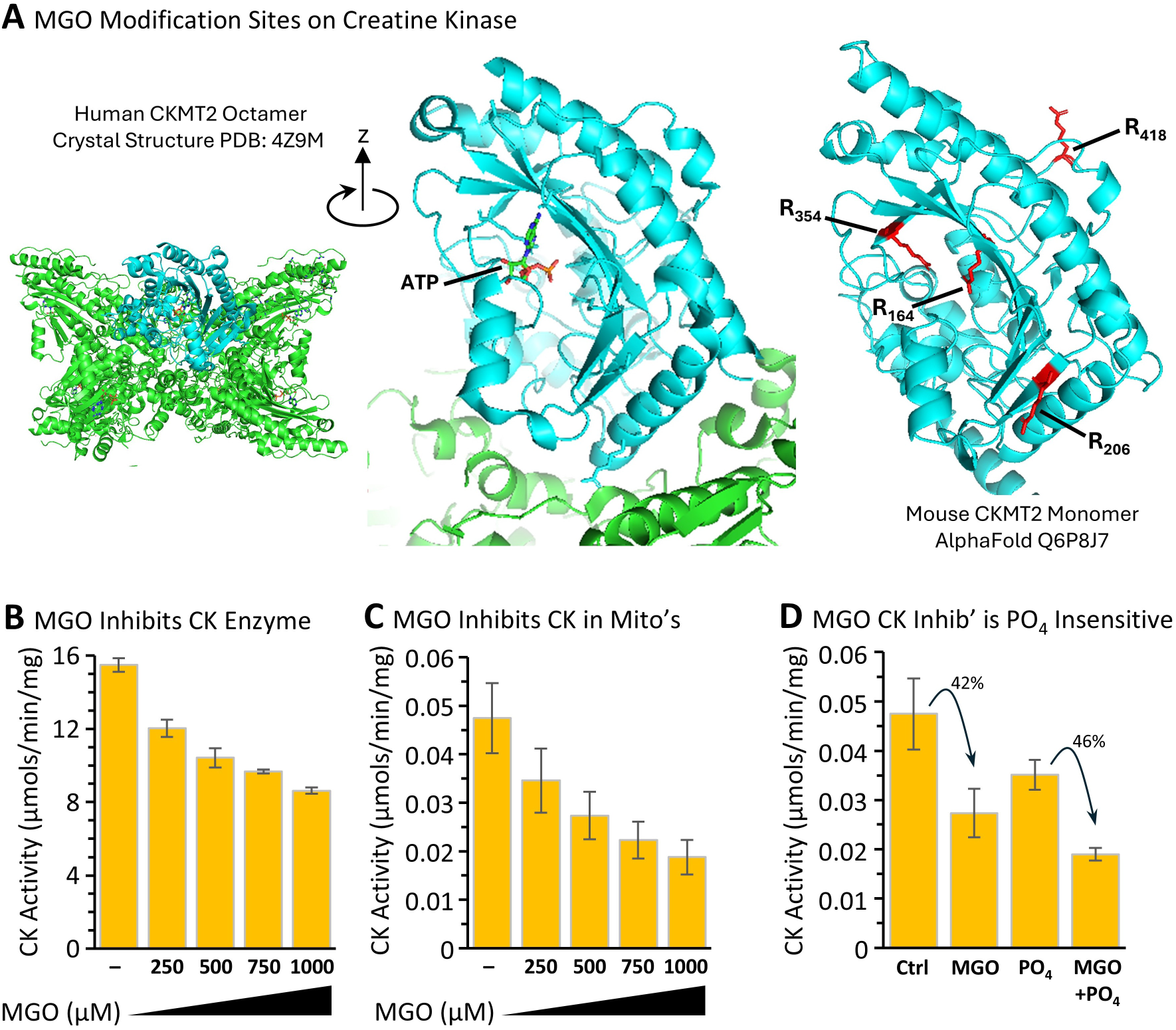
MGO Modification and Inhibition of Creatine Kinase. **(A)**: Left Panel shows the crystal structure of octameric human mitochondrial creatine kinase (CKMT2, pdb-4z9m). One monomer is shown in cyan with the other 7 CK units in green. Central panel shows a human CKMT2 monomer, rotated ∼ 20° clockwise around the z-axis relative to the octamer, highlighting ATP binding within the active site of the enzyme. Right panel shows the AlphaFold predicted structure of mouse CKMT2 (accession #Q6P8J7) in the same approximate orientation as the human monomer, highlighting 4 residues that are modified by MGO in a phosphate sensitive manner (see Fig. 3B). **(B):** Enzyme activity of purified CK, pre-incubated with indicated concentrations of MGO. **(C):** Activity of CK in isolated mitochondria pre-incubated with the indicated concentrations of MGO. **(D):** Activity of CK in mitochondria pre-incubated with 0.5 mM MGO, 3 mM phosphate, or MGO plus phosphate. Percent inhibition by MGO in each condition (±phosphate) is shown. CK activity data are means ± SEM, N=3 independent preparations.

Together with our proteomic observations (**Fig. 3**), these data indicate that the phenomena of CK modification by MGO (which is phosphate sensitive) and CK inhibition by MGO (which is phosphate insensitive) are mechanistically unrelated. This could be due to other residues required for CK activity being targeted by MGO but missed by our proteomic workflow (e.g., C_278_ which is known to be covalently modified by a novel CK inhibitor ^[55]^).

These observations also present a paradox for MGO regulation of the PT pore: MGO’s effect on the PT pore was phosphate sensitive (**Fig. 3A**), while MGO’s effect on mitochondrial CK activity was not (**Fig. 4D**). This suggests that CK activity itself is not important for the PT pore. Indeed, as shown in **Suppl. Fig. 4**, addition of a wide range of concentrations of the CK inhibitor FDNB, including in the presence of phosphate, had no effect on PT pore opening. In addition, a newly described CK inhibitor (CKi-1 ^[55]^) did not impact PT pore opening. This result suggests MGO may indeed regulate CK activity, but this is unrelated to the PT pore. In a similar manner, it has been shown that the inhibitory effect of MGO on mitochondrial respiration can be prevented by high concentrations of creatine but this effect did not depend on CK enzymatic activity ^[56]^.

As **Fig. 4A** shows, there were two other phosphate sensitive glycation sites in CK (R_206_ and R_418_) but these are far away from the active site and their importance for the PT pore effect of MGO is unclear. Five additional MGO modifications in the CK enzyme (R_158_, R_186_, K_276_, R_270_, K_419_) were not impacted by phosphate, so their importance for PT pore opening is unclear.

## Discussion

In this study, the discovery that the effect of MGO on the mitochondrial PT pore was inhibited by phosphate led to the identification of mitochondrial creatine kinase (CK) as a potential mediator of these effects. Although MGO indeed inhibited the enzyme activity of CK, this did not appear to be phosphate sensitive, and CK inhibition alone did not impact the PT pore. Together these findings (**Graphical Abstract Figure**) suggest additional mechanisms may underlie the combined effects of MGO and phosphate on the PT pore.

Previous studies found that mitochondrial CK could be co-purified with ANT and VDAC and reconstituted into membranes, whereupon the breakdown of CK from its octameric to dimeric form revealed a CsA-sensitive permeability similar to the PT pore ^[57,58]^. These experiments, in either brain mitochondria or in liver mitochondria from transgenic mice overexpressing *Ckmt2*, suggest that octameric CK is inhibitory toward the PT pore, while the CK dimer promotes the pore. In turn, this suggests that modifications to CK that impact octamer assembly could be important in regulating PT pore formation. One such modification identified herein is R_206_, at the edge of the β-sheet (**Fig. 4A**). Structurally, this residue appears to assist in coordinating an α-helix that is the interface for multimerization of CK subunits. However, modification of this residue by MGO would be expected to disrupt CK multimer assembly and thus promote pore opening. As such, the inhibitory effect of MGO on the PT pore suggests modification at this residue is not important.

Further relating to CK assembly and structure, notably both R_418_ (phosphate sensitive) and K_419_ (phosphate insensitive) are at the C-terminus of the enzyme. While early work suggested the N-terminus of CK was important for binding to cardiolipin in mitochondrial membranes ^[59]^, it was later discovered that positively charged residues in the C-terminus are also required for this function ^[60]^. As such, MGO modification at the C-terminus would be expected to neutralize such charges, potentially dissociating CK from the membrane and inhibiting its involvement in the PT pore. Clearly, further studies of CK quaternary structure in the presence of MGO and/or phosphate are warranted.

In addition to creatine kinase, numerous other mitochondrial MGO targets were identified that could potentially be relevant to the PT pore. Among these, K_192_ of ATP synthase subunit O (a.k.a. OSCP) may be important, because this protein in the stator/cap of the complex is the binding site for the pore regulator CypD ^[42]^. Several other residues within the ATP synthase complex were also MGO targets, including R_73_, R_347_, R_186_, R_424_, K_427_, R_527_ and K_531_ in the α-subunit (*Atp5f1a*), plus K_350_, R_422_ and K_426_ in the β-subunit (*Atp5f1b*). These modifications may underlie some of the known effects of MGO on mitochondrial respiration ^[56]^. Furthermore ANT1 (*Slc25a4*) was modified at several residues including R_105_, K_107_ and K_260_, The latter is located on a loop facing the mitochondrial intermembrane space, which may be important for interfacing with proteins including CK.

It has not escaped our notice that a common theme pervades many of the mitochondrial MGO target proteins identified herein – they are not only implicated in the PT pore, but they are also all involved in high energy phosphate metabolism (ATP / ADP / Cr / PCr / Pi). Notably a previous study found extensive glycation of cardiomyocyte contractile filaments ^[52,53]^, which consume ATP as a substrate during contractile cycling. Together this suggests that proteins with elevated flux of high energy phosphates may be particularly susceptible to MGO modification.

A limitation of this study was our focus on only CEL and MG-H1 as products of MGO modification on lysine and arginine residues respectively. It is also known that MGO can modify guanosine residues ^[61]^ and can cause chromosomal instability ^[6]^, and can react with other secondary amines (this is the basis for use of the scavenging molecule aminoguanidine). Tryptophan, histidine, asparagine and glutamine may also be susceptible to MGO glycation depending on the steric accessibility of their amines. MGO is also known to drive formation of cross-links between proteins ^[62]^, but the significance of this in mitochondria is not known.

The physiologic relevance of these findings is likely complex. As mentioned in the introduction there are numerous cardioprotective scenarios in which glycolysis is elevated. We showed that the *Alkbh7*^-/-^ mouse is cardioprotected against IR injury and exhibits hyperglycation of cardiac and mitochondrial proteins ^[25]^. Furthermore, as shown in **Suppl. Fig. 1**, three other models of cardioprotection exhibit more glycative stress: (i) ischemic preconditioning, (ii) glucose-insulin-potassium, (iii) the Glyco-Hi mouse heart. Together with the finding that MGO can inhibit the PT pore, these results fundamentally question the dogma that MGO is solely pathologic, and suggest a more nuanced view of this metabolite, in which it may be able to signal cardioprotection. Such a signaling role would position MGO alongside ROS as another example of hormesis, in which low levels of normally toxic species may have beneficial effects.

## Acknowledgements, Funding, Conflict-of-Interest

This work was funded by a grant (to PSB) from the National Institutes of Health (R01-HL071158). RL was funded in part by T32GM135134. We thank Kevin Welle in the URMC Proteomics Core Facility for help with proteomics experiments. We thank Bradford Hill (University of Louisville) for the generous gift of heart samples from wild-type and Glyco-Hi mice. All authors have no commercial, financial, or other conflicts of interest with regard to the content of this paper.

## Data Availability

All data for the figures presented herein are available on the data sharing site FigShare (DOI: 10.6084/m9.figshare.34047288)

## Supplemental Information

**Supplemental Figure 1.**
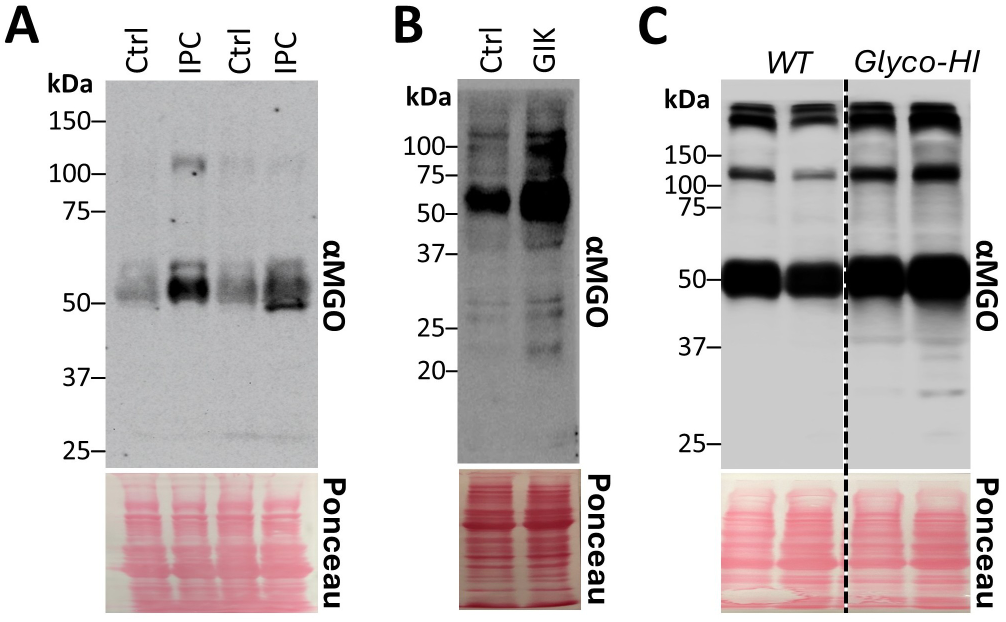
Glycation of myocardial proteins under cardioprotective scenarios. **(A):** Mouse hearts were subjected to control perfusion or IPC as per the methods. **(B):** Mouse hearts were subjected to GIK therapy as per the methods. **(C):** Heart tissue was obtained from wild-type vs. Glyco-Hi mice. Heart tissue was separated by SDS-PAGE then western blotted for anti-MGO protein adducts. Ponceau stained membranes are shown below for protein loading. Blots are representative of at least 3 independent samples per group.

**Supplemental Figure 2.**
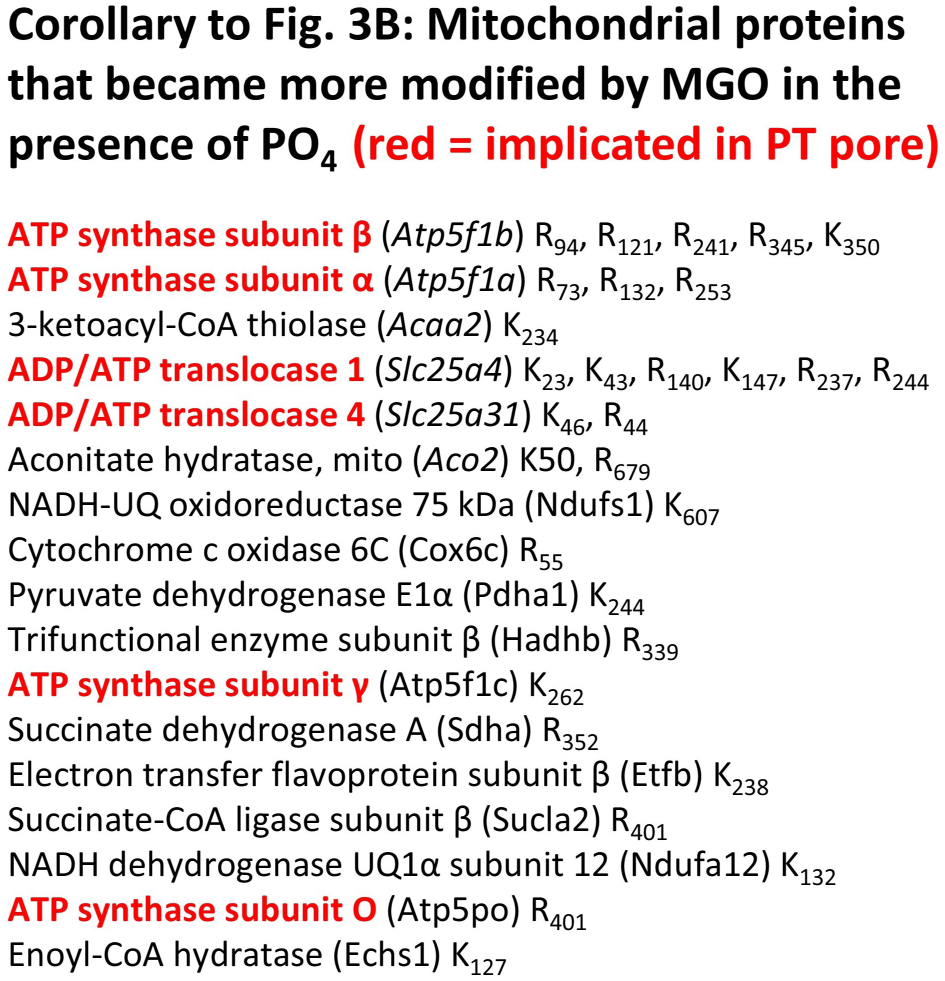
Corollary to Figure 3B. Table shows 30 modified peptides that increased in abundance by >1.5-fold when phosphate was present. Red indicates proteins implicated in PT pore opening.

**Supplemental Figure 3.**
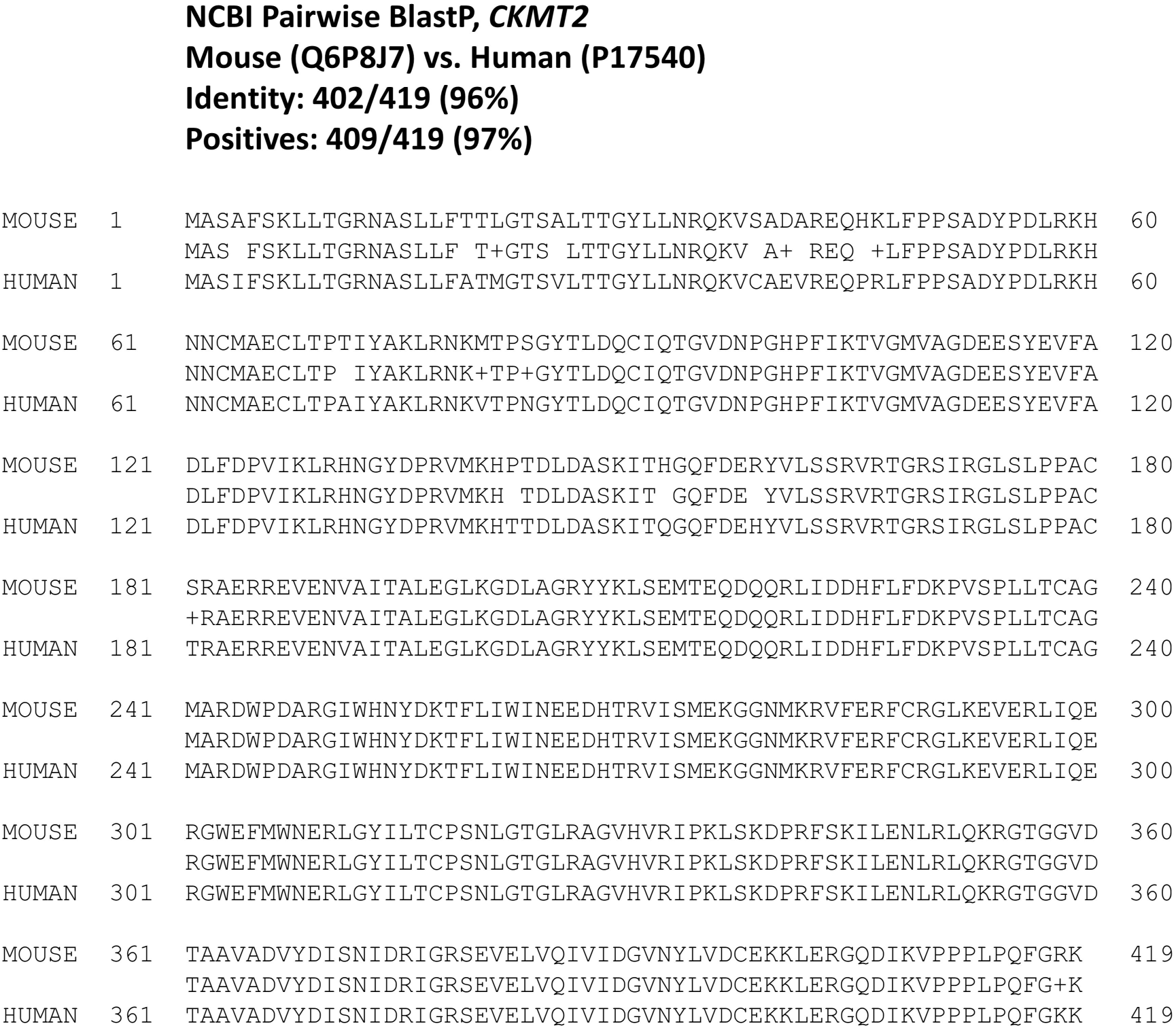
Sequence alignment (NCBI pairwise BlastP) between human CKMT2 (accession #P17450) and mouse CKMT2 (accession #Q6P8J7) thus justifying the use of AlphaFold predicted sequence for structural modeling of MGO modifications in Figure 4A.

**Supplemental Figure 4.**
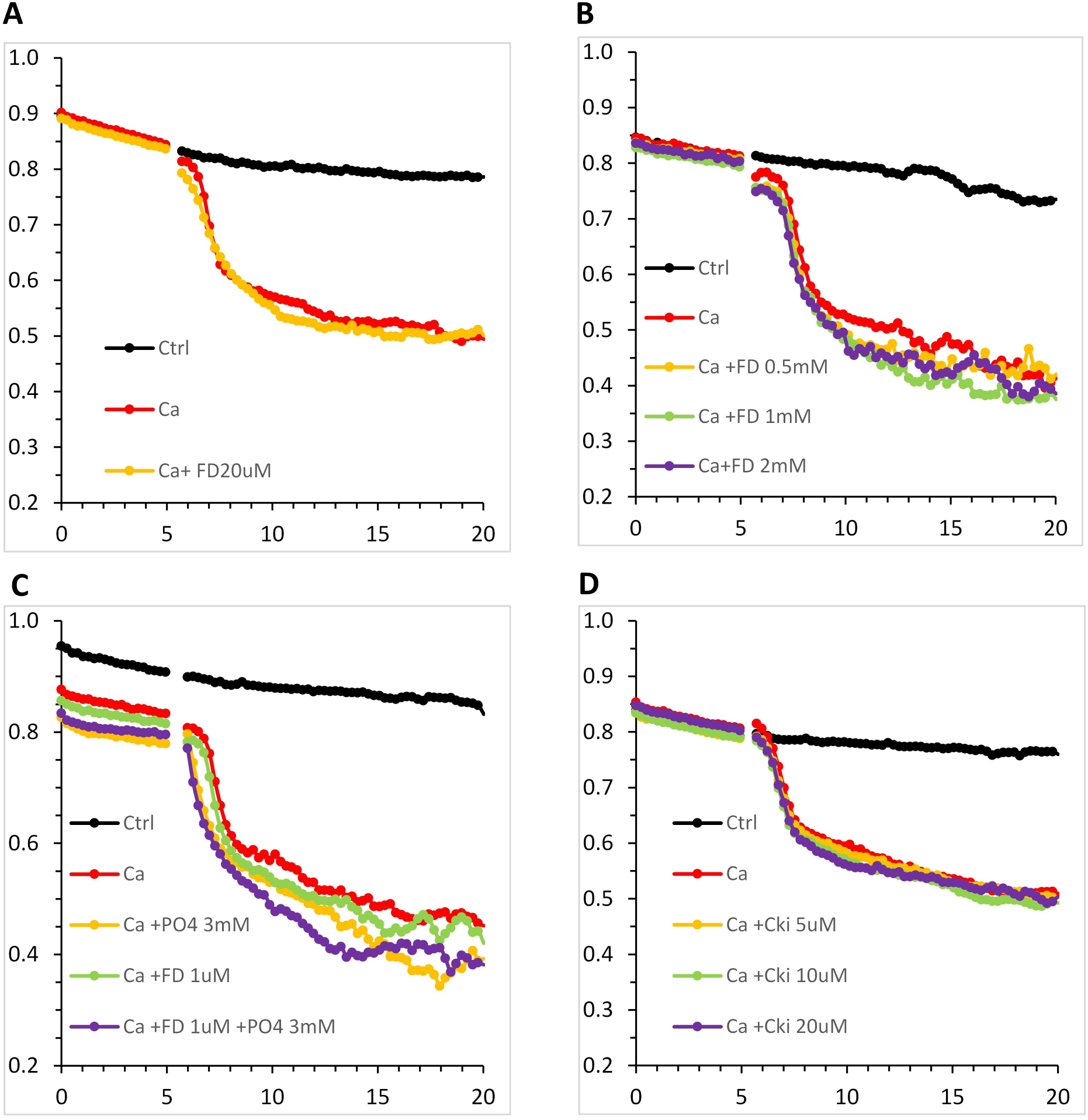
Effect of CK inhibitors on PT Pore Opening. Heart mitochondria were subjected to Ca^2+^ induced PT pore opening as in Fig. 3A. Each panel is an independent mitochondrial preparation **(A):** The CK inhibitor FDNB was tested at 20 μM. **(B):** The CK inhibitor FDNB was tested at 0.5, 1, and 2 mM. **(C):** The CK inhibitor FDNB was tested at 1 μM in the absence or presence of phosphate. **(D):** The novel CK inhibitor CKi-1 was tested at 5, 10, and 20 μM (IC_50_ for the compound is 1.2 μM, Darabedian *et al.* (2023) *Nat Chem Biol* **19**, 815-824).

## Notes

https://figshare.com/articles/dataset/Dataset_for_Lake_et_al_MGO_CK_paper/34047288?file=69545886

### Competing Interest Statement

The authors have declared no competing interest.

## References Cited

1. Walters, A.M., Porter, G.A., Jr. & Brookes, P.S. (2012) Mitochondria as a drug target in ischemic heart disease and cardiomyopathy. Circ Res 111, 1222–36.

2. Di Lisa, F., Canton, M., Menabo, R., Dodoni, G. & Bernardi, P. (2003) Mitochondria and reperfusion injury. The role of permeability transition. Basic Res Cardiol 98, 235–41.

3. Twarda-Clapa, A., Olczak, A., Bialkowska, A.M. & Koziolkiewicz, M. (2022) Advanced Glycation End-Products (AGEs): Formation, Chemistry, Classification, Receptors, and Diseases Related to AGEs. Cells 11.

4. Vaskova, J., Kovacova, G., Pudelsky, J., Palencar, D. & Mickova, H. (2025) Methylglyoxal Formation-Metabolic Routes and Consequences. Antioxidants (Basel) 14.

5. Rabbani, N. & Thornalley, P.J. (2011) Glyoxalase in diabetes, obesity and related disorders. Semin Cell Dev Biol 22, 309–17.

6. Donnellan, L., Simpson, B., Dhillon, V.S., Costabile, M., Fenech, M. & Deo, P. (2021) Methylglyoxal induces chromosomal instability and mitotic dysfunction in lymphocytes. Mutagenesis 36, 339–348.

7. Rabbani, N. & Thornalley, P.J. (2014) Measurement of methylglyoxal by stable isotopic dilution analysis LC-MS/MS with corroborative prediction in physiological samples. Nat Protoc 9, 1969–79.

8. Moon, C.H., Jung, Y.S., Lee, S.H. & Baik, E.J. (1999) Protein kinase C inhibitors abolish the increased resistance of diabetic rat heart to ischemia-reperfusion injury. Jpn J Physiol 49, 409–15.

9. Murarikova, M., Ferko, M., Waczulikova, I., Jasova, M., Kancirova, I., Murinova, J. & Ravingerova, T. (2017) Changes in mitochondrial properties may contribute to enhanced resistance to ischemia-reperfusion injury in the diabetic rat heart. Can J Physiol Pharmacol 95, 969–976.

10. Ravingerova, T., Stetka, R., Volkovova, K., Pancza, D., Dzurba, A., Ziegelhoffer, A. & Styk, J. (2000) Acute diabetes modulates response to ischemia in isolated rat heart. Mol Cell Biochem 210, 143–51.

11. Hanssen, N.M.J., Scheijen, J., Houben, A., van de Waarenburg, M., Berendschot, T., Webers, C.A.B., Reesink, K.D., van Greevenbroek, M.M.J., van der Kallen, C., Schaper, N.C., Schram, M.T., Henry, R.M.A., Stehouwer, C.D.A. & Schalkwijk, C.G. (2021) Fasting and post-oral-glucose-load levels of methylglyoxal are associated with microvascular, but not macrovascular, disease in individuals with and without (pre)diabetes: The Maastricht Study. Diabetes Metab 47, 101148.

12. Maasen, K., Eussen, S., Dagnelie, P.C., Houben, A., Webers, C.A.B., Schram, M.T., Berendschot, T., Stehouwer, C.D.A., Opperhuizen, A., van Greevenbroek, M.M.J. & Schalkwijk, C.G. (2022) Habitual intake of dietary methylglyoxal is associated with less low-grade inflammation: the Maastricht Study. Am J Clin Nutr 116, 1715–1728.

13. Mayen, A.L., Maasen, K., Hana, C., Knaze, V., Scheijen, J., Eussen, S., Vangrieken, P., Debras, C., Blanco, J., Dahm, C.C., Aleksandrova, K., Schulze, M.B., Dansero, L., Masala, G., Panico, S., Sieri, S., Guevara, M., Moreno Iribas, C., Petrova, D., Santiuste, C., Zamora-Ros, R., van der Schouw, Y.T., Aglago, E., Huybrechts, I., Freisling, H., Schalkwijk, C. & Jenab, M. (2025) Higher intakes of dietary dicarbonyl compounds are associated with lower risk of cardiovascular disease. Eur J Prev Cardiol 32, 1588–1600.

14. Berends, E., Vangrieken, P., Amiri, N., van de Waarenburg, M.P.H., Scheijen, J., Hermes, D., Wouters, K., van Oostenbrugge, R.J., Schalkwijk, C.G. & Foulquier, S. (2025) Increased Levels of Circulating Methylglyoxal Have No Consequence for Cerebral Microvascular Integrity and Cognitive Function in Young Healthy Mice. Mol Neurobiol 62, 4190–4202.

15. Van den Eynde, M.D.G., Houben, A., Scheijen, J., Linkens, A.M.A., Niessen, P.M., Simons, N., Hanssen, N.M.J., Kusters, Y., Eussen, S., Miyata, T., Stehouwer, C.D.A. & Schalkwijk, C.G. (2023) Pyridoxamine reduces methylglyoxal and markers of glycation and endothelial dysfunction, but does not improve insulin sensitivity or vascular function in abdominally obese individuals: A randomized double-blind placebo-controlled trial. Diabetes Obes Metab 25, 1280–1291.

16. Zemva, J., Fink, C.A., Fleming, T.H., Schmidt, L., Loft, A., Herzig, S., Kniess, R.A., Mayer, M., Bukau, B., Nawroth, P.P. & Tyedmers, J. (2017) Hormesis enables cells to handle accumulating toxic metabolites during increased energy flux. Redox Biol 13, 674–686.

17. Bollong, M.J., Lee, G., Coukos, J.S., Yun, H., Zambaldo, C., Chang, J.W., Chin, E.N., Ahmad, I., Chatterjee, A.K., Lairson, L.L., Schultz, P.G. & Moellering, R.E. (2018) A metabolite-derived protein modification integrates glycolysis with KEAP1-NRF2 signalling. Nature 562, 600–604.

18. Nadtochiy, S.M., Redman, E., Rahman, I. & Brookes, P.S. (2011) Lysine deacetylation in ischaemic preconditioning: the role of SIRT1. Cardiovasc Res 89, 643–9.

19. Nadtochiy, S.M., Urciuoli, W., Zhang, J., Schafer, X., Munger, J. & Brookes, P.S. (2015) Metabolomic profiling of the heart during acute ischemic preconditioning reveals a role for SIRT1 in rapid cardioprotective metabolic adaptation. J Mol Cell Cardiol 88, 64–72.

20. Wang, Q., Donthi, R.V., Wang, J., Lange, A.J., Watson, L.J., Jones, S.P. & Epstein, P.N. (2008) Cardiac phosphatase-deficient 6-phosphofructo-2-kinase/fructose-2,6-bisphosphatase increases glycolysis, hypertrophy, and myocyte resistance to hypoxia. Am J Physiol Heart Circ Physiol 294, H2889–97.

21. Grossman, A.N., Opie, L.H., Beshansky, J.R., Ingwall, J.S., Rackley, C.E. & Selker, H.P. (2013) Glucose-insulin-potassium revived: current status in acute coronary syndromes and the energy-depleted heart. Circulation 127, 1040–8.

22. Chu, L.M., Osipov, R.M., Robich, M.P., Feng, J., Oyamada, S., Bianchi, C. & Sellke, F.W. (2010) Is hyperglycemia bad for the heart during acute ischemia? J Thorac Cardiovasc Surg 140, 1345–52.

23. Apstein, C.S. & Opie, L.H. (1999) Glucose-insulin-potassium (GIK) for acute myocardial infarction: a negative study with a positive value. Cardiovasc Drugs Ther 13, 185–9.

24. Maroko, P.R., Libby, P., Sobel, B.E., Bloor, C.M., Sybers, H.D., Shell, W.E., Covell, J.W. & Braunwald, E. (1972) Effect of glucose-insulin-potassium infusion on myocardial infarction following experimental coronary artery occlusion. Circulation 45, 1160–75.

25. Kulkarni, C.A., Nadtochiy, S.M., Kennedy, L., Zhang, J., Chhim, S., Alwaseem, H., Murphy, E., Fu, D. & Brookes, P.S. (2020) ALKBH7 mediates necrosis via rewiring of glyoxal metabolism. Elife 9.

26. Nadtochiy, S.M., Wang, Y.T., Nehrke, K., Munger, J. & Brookes, P.S. (2018) Cardioprotection by nicotinamide mononucleotide (NMN): Involvement of glycolysis and acidic pH. J Mol Cell Cardiol 121, 155–162.

27. Miura, T., Tanno, M. & Sato, T. (2010) Mitochondrial kinase signalling pathways in myocardial protection from ischaemia/reperfusion-induced necrosis. Cardiovasc Res 88, 7–15.

28. Solenkova, N.V., Solodushko, V., Cohen, M.V. & Downey, J.M. (2006) Endogenous adenosine protects preconditioned heart during early minutes of reperfusion by activating Akt. Am J Physiol Heart Circ Physiol 290, H441–9.

29. Simm, A., Muller, B., Nass, N., Hofmann, B., Bushnaq, H., Silber, R.E. & Bartling, B. (2015) Protein glycation - Between tissue aging and protection. Exp Gerontol 68, 71–5.

30. Bravo, C., Kudej, R.K., Yuan, C., Yoon, S., Ge, H., Park, J.Y., Tian, B., Stanley, W.C., Vatner, S.F., Vatner, D.E. & Yan, L. (2013) Metabolomic analysis of two different models of delayed preconditioning. J Mol Cell Cardiol 55, 19–26.

31. Geng, J., Zhang, Y., Li, S., Li, S., Wang, J., Wang, H., Aa, J. & Wang, G. (2019) Metabolomic Profiling Reveals That Reprogramming of Cerebral Glucose Metabolism Is Involved in Ischemic Preconditioning-Induced Neuroprotection in a Rodent Model of Ischemic Stroke. J Proteome Res 18, 57–68.

32. Ou, Z., Yang, L., Wu, J., Xu, M., Weng, X. & Xu, G. (2024) Metabolic characteristics of ischaemic preconditioning induced performance improvement in Taekwondo athletes using LC‒MS/MS-based plasma metabolomics. Sci Rep 14, 24609.

33. Liu, J., Zhan, G., Chen, D., Chen, J., Yuan, Z.B., Zhang, E.L., Gao, Y.X., Xu, G., Sun, B.D., Liao, W. & Gao, Y.Q. (2017) UPLC-QTOFMS-based metabolomic analysis of the serum of hypoxic preconditioning mice. Mol Med Rep 16, 6828–6836.

34. Eriksson, O., Fontaine, E., Petronilli, V. & Bernardi, P. (1997) Inhibition of the mitochondrial cyclosporin A-sensitive permeability transition pore by the arginine reagent phenylglyoxal. FEBS Lett 409, 361–4.

35. Guo, L., Carraro, M., Sartori, G., Minervini, G., Eriksson, O., Petronilli, V. & Bernardi, P. (2018) Arginine 107 of yeast ATP synthase subunit g mediates sensitivity of the mitochondrial permeability transition to phenylglyoxal. J Biol Chem 293, 14632–14645.

36. Milliken, A.S., Kulkarni, C.A. & Brookes, P.S. (2020) Acid enhancement of ROS generation by complex-I reverse electron transport is balanced by acid inhibition of complex-II: Relevance for tissue reperfusion injury. Redox Biol 37, 101733.

37. Speer, O., Morkunaite-Haimi, S., Liobikas, J., Franck, M., Hensbo, L., Linder, M.D., Kinnunen, P.K., Wallimann, T. & Eriksson, O. (2003) Rapid suppression of mitochondrial permeability transition by methylglyoxal. Role of reversible arginine modification. J Biol Chem 278, 34757–63.

38. Bernardi, P., Gerle, C., Halestrap, A.P., Jonas, E.A., Karch, J., Mnatsakanyan, N., Pavlov, E., Sheu, S.S. & Soukas, A.A. (2023) Identity, structure, and function of the mitochondrial permeability transition pore: controversies, consensus, recent advances, and future directions. Cell Death Differ 30, 1869–1885.

39. Kokoszka, J.E., Waymire, K.G., Levy, S.E., Sligh, J.E., Cai, J., Jones, D.P., MacGregor, G.R. & Wallace, D.C. (2004) The ADP/ATP translocator is not essential for the mitochondrial permeability transition pore. Nature 427, 461–5.

40. Alavian, K.N., Beutner, G., Lazrove, E., Sacchetti, S., Park, H.A., Licznerski, P., Li, H., Nabili, P., Hockensmith, K., Graham, M., Porter, G.A., Jr. & Jonas, E.A. (2014) An uncoupling channel within the c-subunit ring of the F1FO ATP synthase is the mitochondrial permeability transition pore. Proc Natl Acad Sci U S A 111, 10580–5.

41. Tommasin, L., Carrer, A., Nata, F.B., Frigo, E., Fogolari, F., Lippe, G., Carraro, M. & Bernardi, P. (2025) Adenine nucleotide translocator and ATP synthase cooperate in mediating the mitochondrial permeability transition. J Physiol.

42. Giorgio, V., Fogolari, F., Lippe, G. & Bernardi, P. (2019) OSCP subunit of mitochondrial ATP synthase: role in regulation of enzyme function and of its transition to a pore. Br J Pharmacol 176, 4247–4257.

43. Andreeva, L., Tanveer, A. & Crompton, M. (1995) Evidence for the involvement of a membrane-associated cyclosporin-A-binding protein in the Ca(2+)-activated inner membrane pore of heart mitochondria. Eur J Biochem 230, 1125–32.

44. Hausenloy, D.J., Botker, H.E., Engstrom, T., Erlinge, D., Heusch, G., Ibanez, B., Kloner, R.A., Ovize, M., Yellon, D.M. & Garcia-Dorado, D. (2017) Targeting reperfusion injury in patients with ST-segment elevation myocardial infarction: trials and tribulations. Eur Heart J 38, 935–941.

45. da Silva Ferreira, A.C., Reis, S., Rodrigues, C., Oliveira, C. & de Pinho, P.G. (2007) Simultaneous determination of ketoacids and dicarbonyl compounds, key Maillard intermediates on the generation of aged wine aroma. J Food Sci 72, S314–8.

46. Milliken, A.S., Ciesla, J.H., Nadtochiy, S.M. & Brookes, P.S. (2023) Distinct effects of intracellular vs. extracellular acidic pH on the cardiac metabolome during ischemia and reperfusion. J Mol Cell Cardiol 174, 101–114.

47. Lowry, O.H., Rosebrough, N.J., Farr, A.L. & Randall, R.J. (1951) Protein measurement with the Folin phenol reagent. J Biol Chem 193, 265–75.

48. Brookes, P.S., Salinas, E.P., Darley-Usmar, K., Eiserich, J.P., Freeman, B.A., Darley-Usmar, V.M. & Anderson, P.G. (2000) Concentration-dependent effects of nitric oxide on mitochondrial permeability transition and cytochrome c release. J Biol Chem 275, 20474–9.

49. Nadtochiy, S.M., Nauduri, D., Shimanskaya, T.V., Sagach, V.F. & Brookes, P.S. (2008) Purine release: a protective signaling mechanism of the mitochondrial permeability transition pore in ischemia. Fiziol Zh (1SS4) 54, 5–14.

50. Nadtochiy, S.M., Tompkins, A.J. & Brookes, P.S. (2006) Different mechanisms of mitochondrial proton leak in ischaemia/reperfusion injury and preconditioning: implications for pathology and cardioprotection. Biochem J 395, 611–8.

51. Ge, S.X., Jung, D. & Yao, R. (2020) ShinyGO: a graphical gene-set enrichment tool for animals and plants. Bioinformatics 36, 2628–2629.

52. Papadaki, M., Kampaengsri, T., Barrick, S.K., Campbell, S.G., von Lewinski, D., Rainer, P.P., Harris, S.P., Greenberg, M.J. & Kirk, J.A. (2022) Myofilament glycation in diabetes reduces contractility by inhibiting tropomyosin movement, is rescued by cMyBPC domains. J Mol Cell Cardiol 162, 1–9.

53. Delligatti, C.E. & Kirk, J.A. (2024) Glycation in the cardiomyocyte. Vitam Horm 125, 47–88.

54. Hall, N. & DeLuca, M. (1984) The effect of inorganic phosphate on creatine kinase in respiring rat heart mitochondria. Arch Biochem Biophys 229, 477–82.

55. Darabedian, N., Ji, W., Fan, M., Lin, S., Seo, H.S., Vinogradova, E.V., Yaron, T.M., Mills, E.L., Xiao, H., Senkane, K., Huntsman, E.M., Johnson, J.L., Che, J., Cantley, L.C., Cravatt, B.F., Dhe-Paganon, S., Stegmaier, K., Zhang, T., Gray, N.S. & Chouchani, E.T. (2023) Depletion of creatine phosphagen energetics with a covalent creatine kinase inhibitor. Nat Chem Biol 19, 815–824.

56. Roy, S.S., Biswas, S., Ray, M. & Ray, S. (2003) Protective effect of creatine against inhibition by methylglyoxal of mitochondrial respiration of cardiac cells. Biochem J 372, 661–9.

57. O’Gorman, E., Beutner, G., Dolder, M., Koretsky, A.P., Brdiczka, D. & Wallimann, T. (1997) The role of creatine kinase in inhibition of mitochondrial permeability transition. FEBS Lett 414, 253–7.

58. Beutner, G., Ruck, A., Riede, B., Welte, W. & Brdiczka, D. (1996) Complexes between kinases, mitochondrial porin and adenylate translocator in rat brain resemble the permeability transition pore. FEBS Lett 396, 189–95.

59. Cheneval, D. & Carafoli, E. (1988) Identification and primary structure of the cardiolipin-binding domain of mitochondrial creatine kinase. Eur J Biochem 171, 1–9.

60. Schlattner, U., Gehring, F., Vernoux, N., Tokarska-Schlattner, M., Neumann, D., Marcillat, O., Vial, C. & Wallimann, T. (2004) C-terminal lysines determine phospholipid interaction of sarcomeric mitochondrial creatine kinase. J Biol Chem 279, 24334–42.

61. Mazza, M.C., Shuck, S.C., Lin, J., Moxley, M.A., Termini, J., Cookson, M.R. & Wilson, M.A. (2022) DJ-1 is not a deglycase and makes a modest contribution to cellular defense against methylglyoxal damage in neurons. J Neurochem 162, 245–261.

62. Mitra, B., Lametsch, R., Greco, I. & Ruiz-Carrascal, J. (2018) Advanced glycation end products, protein crosslinks and post translational modifications in pork subjected to different heat treatments. Meat Sci 145, 415–424.

